# Embodied virtual reality for the study of real-world motor learning

**DOI:** 10.1101/2020.03.19.998476

**Authors:** Shlomi Haar, Guhan Sundar, A. Aldo Faisal

**Affiliations:** Brain and Behaviour Lab: Dept. of Bioengineering, Imperial College London, London, UK; Dept. of Computing, Imperial College London, London, UK; UKRI Centre for Doctoral Training in AI for Healthcare, Imperial College London, London, UK; MRC London Institute of Medical Sciences, Imperial College London, London, UK

**Keywords:** motor learning, motor skill, real-world, full-body movement, virtual reality, embodiment, motor neuroscience

## Abstract

Motor-learning literature focuses on simple laboratory-tasks due to their controlled manner and the ease to apply manipulations to induce learning and adaptation. Recently, we introduced a billiards paradigm and demonstrated the feasibility of real-world-neuroscience using wearables for naturalistic full-body motion-tracking and mobile-brain-imaging. Here we developed an embodied virtual-reality (VR) environment to our real-world billiards paradigm, which allows to control the visual feedback for this complex real-world task, while maintaining sense of embodiment. The setup was validated by comparing real-world ball trajectories with the trajectories of the virtual balls, calculated by the physics engine. We then ran our learning protocol in the embodied VR. Subjects played billiard shots when they held the physical cue and hit a physical ball on the table while seeing it all in VR. We found comparable learning trends in the embodied VR to those we previously reported in the physical real-world task. Embodied VR can be used for learning real-world tasks in a highly controlled environment which enables applying visual manipulations, common in laboratory-tasks and rehabilitation, to a real-world full-body task. Embodied VR enables to manipulate feedback and apply perturbations to isolate and assess interactions between specific motor-learning components, thus enabling addressing the current questions of motor-learning in real-world tasks. Such a setup can be used for rehabilitation, where VR is gaining popularity but the transfer to the real-world is currently limited, presumably, due to the lack of embodiment.

## Introduction

Motor learning is a key feature of our development and our daily lives, from a baby learning to crawl, to an adult learning crafts or sports, or undergoing rehabilitation after an injury or a stroke. It is a complex process, which involves movement in many degrees of freedom (DoF) and multiple learning mechanisms. Yet the majority of motor learning literature focuses on simple lab-based tasks with limited DoF. The key advantage of these tasks (which made them so popular) is the ability to apply highly controlled manipulations. These manipulations can be a haptic perturbation, where the robotic manipulandum pushes the subjects in a different direction from their intended movement, such as force-field adaptation [e.g. 1–4]. Alternatively, it can be a visual manipulation where the object that represents the subject’s end effector is moving in a different direction, speed, or magnitude than the end effector itself, such as in visuomotor rotation adaptation [e.g. 5–9]. Additionally, visual manipulations can be applied to the feedback, by adding delays [10–13], or showing online feedback of the full movement trajectory versus only the end-point for knowledge of results [e.g. 14–16]. These manipulations allow to isolate specific movement/learning components and establish causality.

In contrast to lab-based tasks, real-world neuroscience approaches study neurobehavioral processes in natural behavioural settings [17–20]. We recently presented a naturalistic real-world motor learning paradigm, using wearables for full-body motion tracking and EEG for mobile brain imaging, while making people perform actual real-world tasks, such as playing the competitive sport of pool-table billiards [21,22]. We showed that motor learning is a full-body process that involves multiple learning mechanisms, and different subjects might prefer one over the other. While the study of real-world tasks takes us closer to understanding real-world motor-learning, it is lacking the key advantage of lab-based toy-tasks, highly controlled manipulations of known variables.

To mechanistically study the human brain and cognition in real-world tasks we have to be able to introduce causal manipulations. This motivated us now to develop and evaluate a real-world motor learning paradigm using a novel experimental framework: Using Virtual Reality (VR [23, see review 24]) to apply controlled visual manipulations in a real-world task. VR has clear benefits such as ease of controlling repetition, feedback, and motivation, as well as overall advantages in safety, time, space, equipment, cost efficiency, and ease of documentation [25,26]. Thus, it is commonly used in rehabilitation after stroke [27,28] or brain injury [29,30], and for Parkinson’s disease [31,32]. In simple sensorimotor lab-based motor learning paradigms, VR training showed to have equivalent results to those of real training [33–35], though adaption in VR appears to be more reliant on explicit/cognitive strategies [35].

While VR is good for visual immersion, it is often lacking the Sense of Embodiment – the senses associated with being inside, having, and controlling a body [36]. Sense of embodiment requires a sense of self-location, agency, and body ownership [37–39]. This study aims to set and validate an Embodied Virtual Reality (EVR) for real-world motor-learning. I.e. VR environment in which all the objects the subjects see and interact with are the physical objects that they can physically sense. This is following the operational definition of embodiment through behaviour which is the ability to process sensorimotor information through technology in the same way as the properties of one’s own body parts [40]. Such EVR setup would enable to apply highly controlled manipulations in a real-world task. We develop here an EVR to our billiards paradigm [21] by synchronizing the positions of the real-world billiards objects (table, cue-stick, balls) into the VR environment using optical marker based motion capture. To be clear, our virtual reality environment was presented simultaneously and veridically matched to the real-world environment, a user looking at the pool table holding a cue, would thus see the pool table and the cue at the same location in the visual field if their VR headset is put on or taken off. Thus, the participant can play with a physical cue and a physical ball on the physical pool-table while seeing the world from the same perspective in VR (see Supplemental Video or https://youtu.be/m68_UYkMbSk). We ran our real-world billiards experimental protocol in this novel EVR to explore the similarities and differences in learning between the real-world paradigm and its EVR mock-up.

## Methods

### Experimental Setup

Our EVR experimental setup (Figure 1A) merges a virtual environment and real-world environment which are presented simultaneously and veridically aligned. It is composed of a real-world environment of a physical pool table, a VR environment of the *same* (Figure 1B), and optical motion capture systems to link between the two environments (Figure 1C). The positions of the virtual billiards table and balls the subjects saw in VR were matched with their respective real-world positions during calibration, and the cue-stick trajectory was streamed into the VR. This allowed subjects to physically engage in the VR task and see their real-world actions and interactions with ball and pool cue stick veridically aligned in VR. This setup enables to apply experimental trickery (which is common in lab-based task but usually impossible in the real-world) to a real-world task (e.g. scaling, up or down, the ball velocity or the directional error to target, adding delay, or hiding the ball’s trajectory).

**Figure 1.**
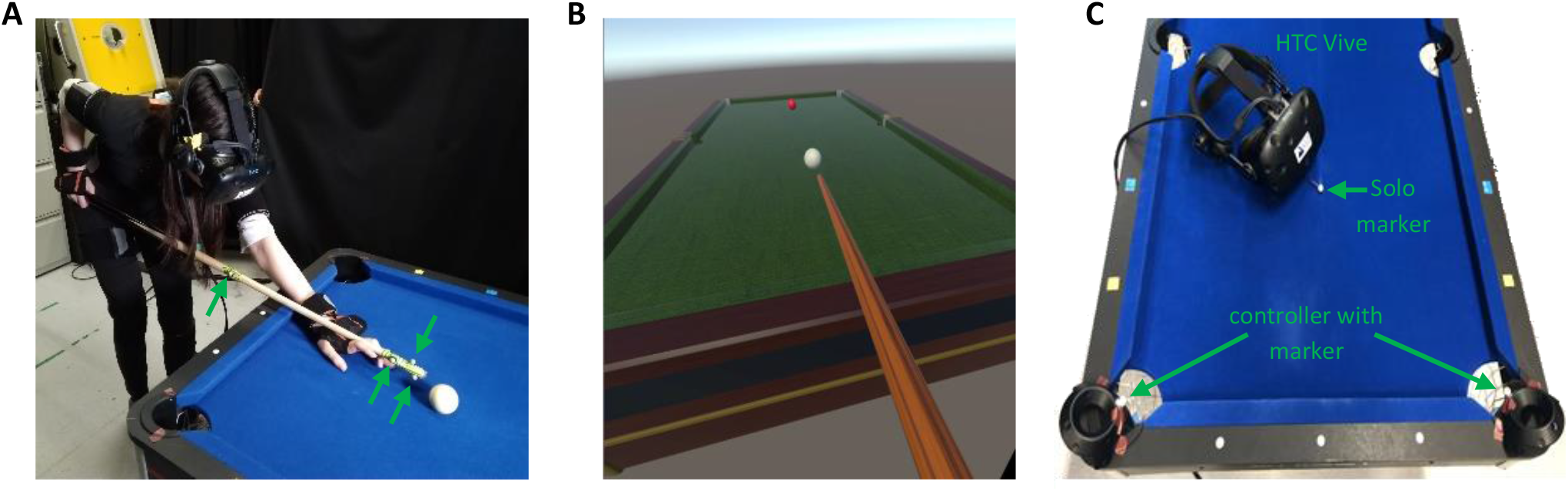
Experimental setup and calibration. (**A**) 10 right-handed healthy subjects performed 300 repeated trials of billiards shots in Embodied Virtual Reality (EVR). Green arrows mark the motion capture markers used to track and stream the cue stick movement into the EVR environment (**B**) Scene view in the EVR. Subjects were instructed to hit the cue ball (white), which was a physical ball on the table (in A), in attempt to shoot the virtual target ball (red) towards the far-left corner. (**C**) For environments calibration, MoCap markers were attached to the HTC Vive controllers which were placed in the pool-table’s pockets with additional solo marker in the cue ball position.

Real-world objects included the same billiards table, cue ball, target ball, and cue stick, used in our real-world billiard study [21]. Subjects were unable to see anything in the real-world environment; they could only see a virtual projection of the game objects. They were, however, able to receive tactile feedback from the objects by interacting with them.

The Optitrack system with 4 Motion Capture cameras (Prime 13W) and Motiv software were used (all made by Natural Point Inc., Corvallis, OR) to stream the position of the real-world cue stick into the VR using 4 markers on the pool cue stick (Figure 1A). The position of each marker was streamed to Unity3d using the NATNET Optitrack Unity3d Client plugin and associated Optitrack Streaming Client script edited for the application. The positions were transformed from the Optitrack environment to the Unity3d environment with a transformation matrix derived during calibration. The cue stick asset was then reconstructed in VR using known geometric quantities of the cue stick and marker locations (Figure 1B). The placement of markers on the cue stick, as well as the position and orientation of the cameras, were key to provide consistent marker tracking and accurate control in VR without significantly constraining the subject movement. The rotation of the cue stick or the position of the subject can interfere with the line of sight between the markers on the cue stick and the cameras. Thus, to prevent errors in cue tracking, if markers become untracked the cue stick disappears from the visual scene until proper tracking is resumed.

The VR billiards environment was built with the Unity3d physics engine (Unity Technologies, San Francisco, CA). The head-mounted display used was the HTC Vive Pro (HTC, Xindian, Taiwan). The frame rate for the VR display was 90 Hz. The Unity3d assets (billiards table, cue stick, balls) were taken from an open-source Unity3d project [41] and scaled to match the dimensions of the real-world objects. Scripts developed in C# to manage game object interactions, apply physics, and record data. Unity3d software was used to develop custom physics for game collisions. Cue stick – cue ball collision force in Unity3d is computed from the median velocity and direction of the cue stick in the 10 frames (~0.11 seconds) before contact. Sensory and auditory feedback comes from the real-world objects for this initial collision. Cue ball – target ball collision is hardcoded as a perfectly inelastic collision. Billiard ball sound effect is outputted to the Vive headphones during this collision. The default Unity3d engine was used for ball dynamics, with specific mass and friction parameters tuned to match as closely as possible to real-world ball behaviour. For the game physics validation, the physical cue ball on the pool table was tracked with a high-speed camera (Dalsa Genie Nano, Teledyne DALSA, Waterloo, Ontario) and its trajectories were compared with those of the VR ball.

For environment calibration, the ‘y-axis’ was set directly upwards (orthogonal to the ground plane) in both the Unity3d and Optitrack environments during their respective initial calibrations. This allows us to only require a 2D (x-z) transformation between environments, using a linear ratio to scale the height. The transformation matrix was determined by matching the positions of known coordinates in both Unity3d and Optitrack environments. We attached markers to the Vive controllers and during calibration mode set them in the corner pockets of the table and placed a solo marker on the cue ball location (Figure 1C), to compute the transformation matrix as well as position and scale of the real-world table. This transformation matrix was then used to transform points from the Optitrack environment into the Unity3d space.

### Billiard ball tracking

We validate the reliability of the trajectories of the virtual balls in the EVR, by comparing them with the trajectories of real balls during the same shot. The movement of the real balls on the pool table were tracked with a computer vision system mounted from the ceiling (Genie Nano C1280 Color Camera, Teledyne Dalsa, Waterloo, Canada), with a resolution of 752×444 pixels and a frequency of 200Hz. Image videos were recorded and analysed with our custom software written for the real-world paradigm [21,22].

### Experimental Design

10 right-handed healthy human volunteers with normal or corrected-to-normal visual acuity (4 women and 6 men, aged 24±2) participated in the study following the experimental protocol from Haar et al [21]. All experimental procedures were approved by the Imperial College Research Ethics Committee and performed in accordance with the declaration of Helsinki. All volunteers gave informed consent prior to participating in the study. The volunteers, who had little to no previous experience with playing billiards, performed 300 repeated trials in the EVR setup where the cue ball (white) and the target ball (red) were placed in the same locations and the subject was asked to shoot the target ball towards the pocket of the far-left corner (Figure 1B). VR trials ended when ball velocities fell below the threshold value, and the next trial began when the subject moved the cue stick tip to a set distance away from the cue ball start position. The trials were split into 6 sets of 50 trials with a short break in-between. For the data analysis, we further split each set into two blocks of 25 trials each, resulting in 12 blocks. During the entire learning process, we recorded the subjects’ full-body movements with a motion-tracking ‘suit’ of 17 wireless inertial measurement units (IMUs). Movement of all game objects in Unity3d (most notably ball and cue stick trajectories relative to the table) were captured in every frame in 90Hz sampling.

### Full-Body Motion Tracking

Kinematic data were recorded at 60 Hz using a wearable motion tracking ‘suit’ of 17 wireless IMUs (Xsens MVN Awinda, Xsens Technologies BV, Enschede, The Netherlands). Data acquisition was done via a graphical interface (MVN Analyze, Xsens Technologies BV, Ensched, The Netherlands). The Xsens joint angles and position data were exported as XML files and analyzed using custom software written in MATLAB (R2017a, The MathWorks, Inc., MA, USA). The Xsens full-body kinematics were extracted in joint angles in 3 degrees of freedom for each joint that followed the International Society of Biomechanics (ISB) recommendations for Euler angle extractions of Z (flexion/extension), X (abduction/adduction) Y (internal/external rotation).

### Analysis of Movement Velocity Profiles

From the sensor data we extracted the joint angular velocity profiles of all joints in all trials. We analysed the joint angular velocity profiles instead of the absolute joint angle’s probability distributions, as the latter are more sensitive to drift in the sensors. We previously showed that joint angular velocity probability distributions are more subject invariant than joint angle distributions suggesting these are the reproducible features across subjects in natural behaviour [42]. In the current task, this robustness is quite intuitive: all subjects stood in front of the same pool table and used the same cue stick, thus the subjects’ body size influenced their joint angles distributions (taller subjects with longer arms had to bend more towards the table and flex their elbow less than shorter subjects with shorter limbs) but not joints angular velocity probability distributions [21]. We defined the peak of the trial as the peak of the average absolute velocity across the DoFs of the right shoulder and the right elbow. We aligned all trials around the peak of the trial and cropped a window of 1 sec around the peak for the analysis of joint angles and velocity profiles.

### Task performance & learning measures

The task performance was measured by the trial error which was defined as an absolute angular difference between the target ball movement vector direction and the desired direction to land the target ball in the centre of the pocket. The decay of error over trials is the clearest signature of learning in the task. To calculate success rates and intertrial variability, the trials were divided into blocks of 25 trials each (each experimental set of 50 trials was divided into two blocks to increase resolution). Success rate in each block was defined by the ratio of successful trial (in which the ball fell into the pocket). To improve robustness and account for outliers, we fitted the errors in each block with a t-distribution and used the location and scale parameters (μ and σ) as the blocks’ centre and variability measures. To correct for learning which is happening within a block, we also calculated a corrected intertrial variability [21], which was the intertrial variability over the residuals from a regression line fitted to the ball direction in each block.

To quantify the within-trial variability structure of the body movement, we use the generalised variance, which is the determinant of the covariance matrix [43] and is intuitively related to the multidimensional scatter of data points around their mean. We measured the generalised variance over the velocity profiles of all joints in each trial to see how it changes with learning in the EVR task relative to the real-world task [21]. To study the complexity of the body movement which was defined by the number of degrees of freedom used by the subject we applied principal component analysis (PCA) across joints for the velocity profiles per trial for each subject and used the number of PCs that explain more than 1% of the variance to quantify the degrees of freedom in each trial movement [21]. We also calculated the manipulative complexity which was suggested by Belić and Faisal [44] as a way to quantify complexity for a given number of PCs on a fixed scale (C = 1 implies that all PCs contribute equally, and C = 0 if one PC explains all data variability).

As a measure of task performance in body space, correlation distances (one minus Pearson correlation coefficient) were calculated between the velocity profile of each joint in each trial to the velocity profiles of that joint in all successful trials. The minimum over these correlation distances produced a single measure of Velocity Profile Error (VPE) for each joint in each trial [21]. While there are multiple combinations of body variables that can all lead to successful task performance, this measure looks for the distance from the nearest successful solution used by the subjects and thus provides a metric that accounts for the redundancy in the body.

## Results

Our embodied Virtual Reality (EVR) framework is presented simultaneously and veridically accurately tracks the real-world environment. We found that players were reliably able to shoot (and aim) the physical pool ball with the physical cue, while their head and eyes were covered and saw the physical scene rendered in virtual reality. We describe in the following, first, how we carried out a validation of the virtual vs physical reality in terms of pool playing task performance and then, report the results of the learning experiments.

### Ball trajectories validation

To validate how well the billiards shot in the EVR resembles the same shot in real life, the cue ball trajectories of 100 shots in various directions (−50°< ø < 50° when 0 is straight forward) were compared between the two environments. The cue ball angles were perfectly correlated (Pearson correlation r=0.99) and the root mean squared error (RMSE) was below 3 degrees (RMSE = 2.85). Thus, the angle of the virtual ball in the EVR, which defines the performance in this billiards task, was very consistent with the angle of the real-world ball (Figure 2A). The velocities were also highly correlated (Pearson correlation r=0.83) between the environments but the ball velocities in the VR were slightly slower than on the real pool-table (Figure 2B), leading to an RMSE of 1.03 m/s.

**Figure 2.**
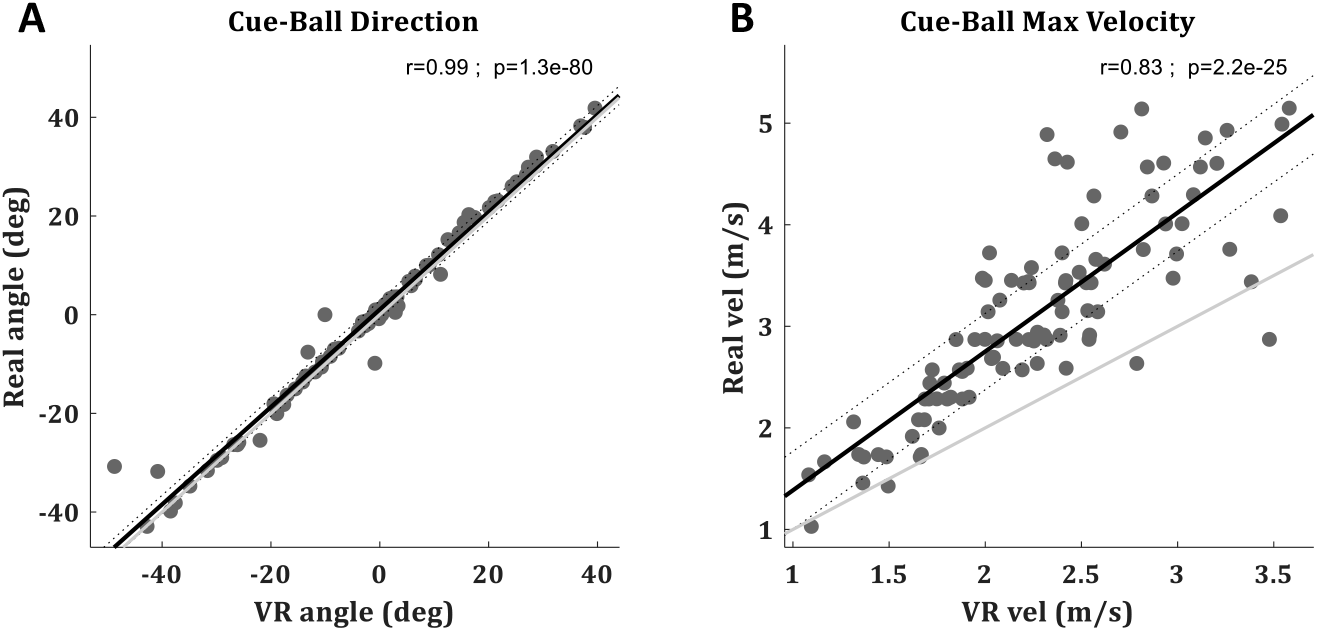
Ball trajectories validation. Within shot comparison between real cue ball trajectory (measured with a high-speed camera) and the virtual ball trajectory (calculated by the VR physics engine). Each one is a shot. A total of 100 billiard shots at various directions (−50°< ø < 50° when 0 is straight forward) are presented. (**A**) Cue ball angles. **(B)** Max velocity of the cue ball during each trial. The regression line is in black with its 95% CI in doted lines. Identity line is in light gray.

### Motor Learning Experiment

To compare the learning of the billiards shot in our EVR to the learning in real-life, we ran the same experimental protocol as in [21] and compared the mean subjects’ performance (Figure 3). In Figure 3, dashed lines are taken from our non-VR physical real-world study [21] which used the identical behaviour protocol, lab environments, table, tracking methods etc, and solid lines are the EVR data from this work. The data of the physical real-world experiment is presented to ease a qualitative comparison between the environments, and thus avoided statistical comparison between the two. Following [21], the trials were divided into blocks of 25 trials each (each experimental set of 50 trials was divided into two blocks to increase resolution) to assess the performance. Over blocks, there is a gradual decay in the mean directional absolute error (Figure 3A). While this decay in the EVR is slower and smaller than in the real-world task, over the session subjects did reduce their error by 8.3±2.47 degrees (mean±SD) and this decay was significant (paired t-test p=0.008). Accordingly, success rates were increasing over blocks (Figure 3B). The success rates increase in the EVR was also lower than in the real-world task. Over the session, subjects improved their success rates by 6.2%± 1.78% (mean±SEM) and this increase was significant (paired t-test p=0.007).

**Figure 3.**
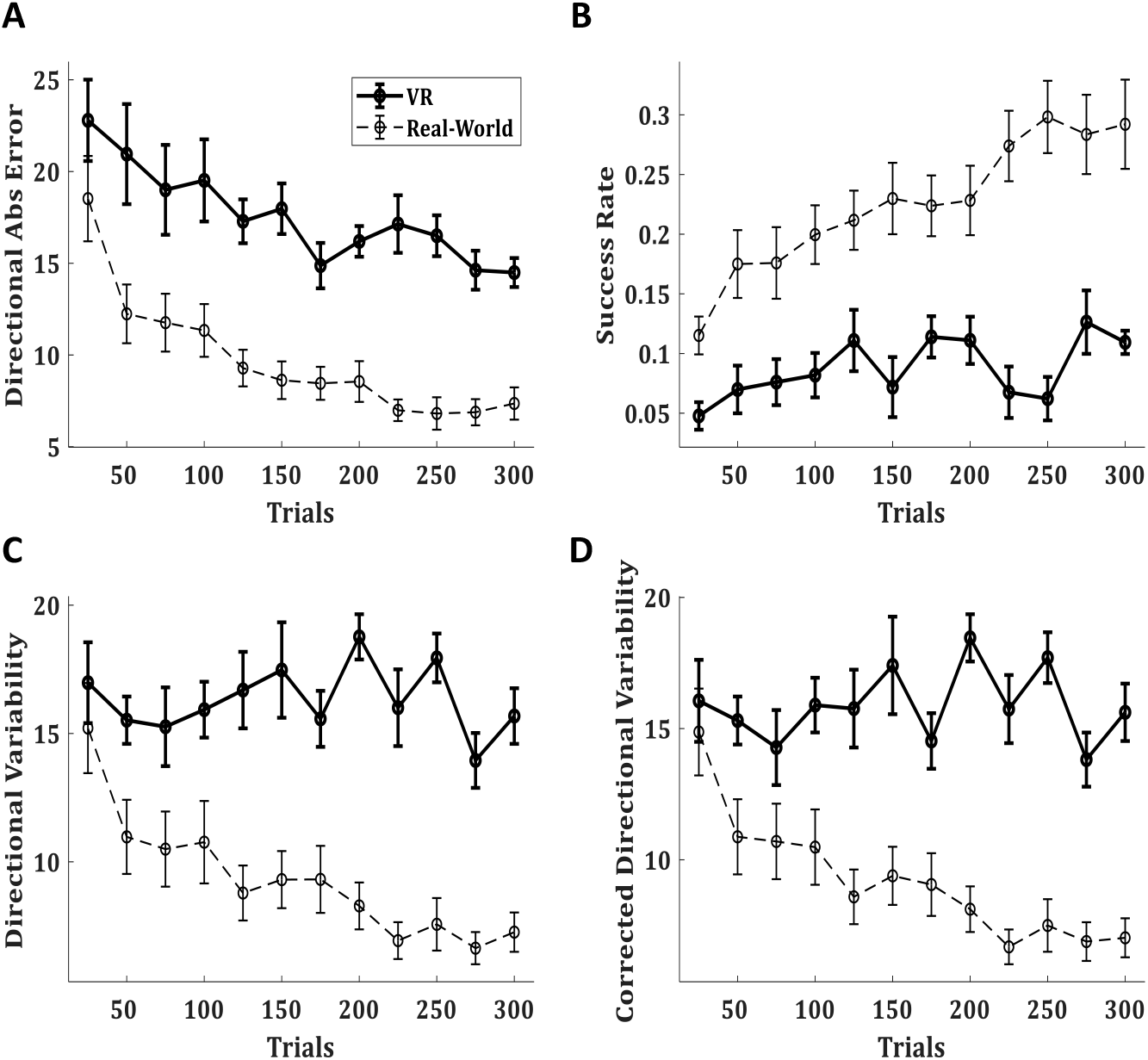
Task performance in EVR vs Real-world. (**A)** The mean absolute directional error of the target-ball, (**B**) The success rate, (**C**) directional variability, and (**D**) directional variability corrected for learning (see text). (**A-D**) presented over blocks of 25 trials. Solid lines present the performance of 10 subjects in the novel EVR environment. For compression, dashed lines present the performance of a group of 30 subjects in the same pool paradigm in the real-world (with no VR) from our previous study [21].

We also see a decay in inter-subject variability over learning, represented by the decrease in the size of the error bar of the directional error over time (Figure 3A). These learning trends in the directional error and success rates are similar to those reported in the real world. Nevertheless, there are clear differences in the learning curve. In the EVR learning occurs slower than in the real-world task, and subjects’ performance is worse. The most striking difference between the environments is in the intertrial variability (Figure 3C). In the real-world task, there was a clear decay in intertrial variability throughout the experiment, whereas in EVR we see no clear trend. Corrected intertrial variability (Figure 3D), calculated to correct for learning happening within the block [21], also showed no learning trend.

The full-body movements were analysed over the velocity profiles of all joints, as those are less sensitive to potential drifts in the IMUs and more robust and reproducible in natural behaviour across subjects and trials [21,42]. The velocity profiles of the different joints in the EVR showed that the movement is in the right arm, as expected. The velocity profiles of the right arm showed the same changes following learning as in the real-world task. The shoulder velocities showed a decrease from the initial trials to the trials of the learning plateau, suggesting less shoulder movement; while the elbow rotation shows an increase in velocity over learning (Figure 4). The covariance matrix over the velocity profiles of the different joints, averaged across blocks of trials of all subjects, emphasizes this trend. Over the first block it shows that most of the variance in the movement is in the right shoulder while in the 9th block (trials 201-225, the beginning of the learning plateau) there is an overall similar structure of the covariance matrix, but with a strong decrease in the shoulder variance and a strong increase in the variance of right elbow rotation (Figure 5A). This is a similar trend to the one observed in the real-world task, and even more robust.

**Figure 4.**
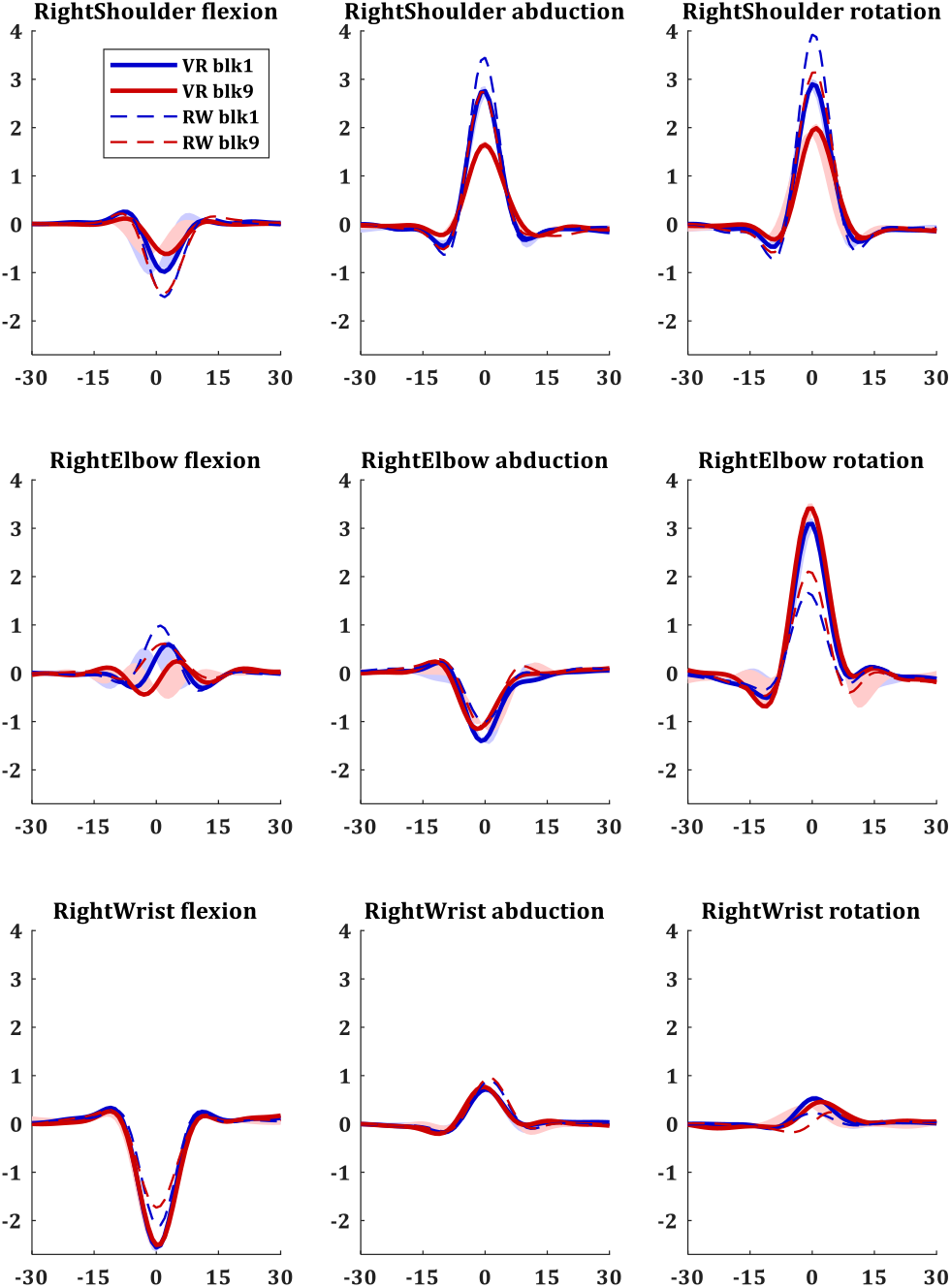
Velocity profiles in EVR vs Real-world. Velocity profiles in 3 degrees of freedom (DoF) for each joint of the right arm joints. Blue lines are the profiles during the first block (trials 1-25), and red lines are the velocity profiles after learning plateaus, during the ninth block (trials 201225). Solid lines present the velocity profiles of 10 subjects in the novel EVR environment. For compression, dashed lines present the velocity profiles of a group of 30 subjects in the same pool paradigm in the real-world (with no VR) from our previous study [21].

**Figure 5.**
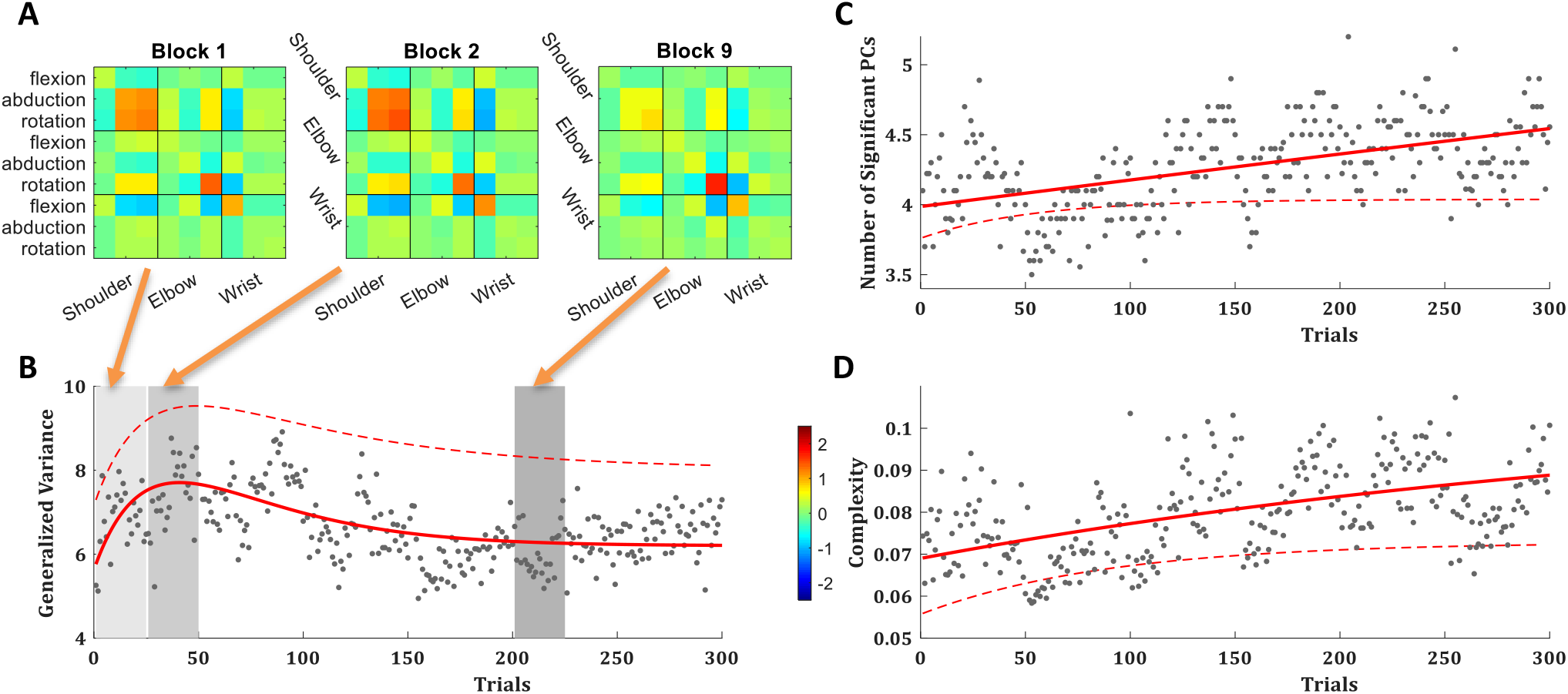
Variance and Complexity comparison. (**A**) The variance covariance matrix of the right arm joints velocity profiles in EVR, averaged across subjects and trials over the first block, second block and the ninth block (after learning plateaus). (**B**) The trial-by-trial generalized variance (GV), with a double-exponential fit (red curve). (**C**) The number of principal components (PCs) that explain more than 1% of the variance in the velocity profiles of all joints in a single trial, with an exponential fit (red curve). (**D**) The manipulative complexity (Belić and Faisal, 2015), with an exponential fit (red curve). (**B-D**) Averaged across all subjects over all trials. Data is averaged over the performance of 10 subjects in the novel EVR environment. Grey dots are the trial averages for the EVR data. Solid red lines are curve fits for EVR data. For compression, dashed lines present the curve fits to a group of 30 subjects in the same pool paradigm in the real-world (with no VR) from our previous study [21].

The generalized variance (GV; the determinant of the covariance matrix [43]) over the velocity profiles of all joints was lower in the EVR than in real-world but showed the same trend: increase fast over the first ~30 trials and later decreased slowly (Figure 5B), suggesting active control of the exploration-exploitation trade-off. The covariance (Figure 5A) shows that the changes in the GV were driven by an initial increase followed by a decrease in the variance of the right shoulder. Like in the real-world, in the EVR as well the internal/external rotation of the right elbow showed a continuous increase in its variance, which did not follow the trend of the GV.

Principal component analysis (PCA) across joints for the velocity profiles per trial for each subject showed that in the EVR subjects used more degrees of freedom in their movement than in the real-world task (Figure 5C&D). While in both environments, in all trials, ~90% of the variance can be explained by the first PC, there is a slow but consistent rise in the number of PCs that explain more than 1% of the variance in the joint velocity profiles (Figure 5C). The manipulative complexity, suggested by Belić and Faisal [44] as a way to quantify complexity for a given number of PCs on a fixed scale (C = 1 implies that all PCs contribute equally, and C = 0 if one PC explains all data variability), showed the same trend (Figure 5D). This suggests that, in both environments, over trials subjects use more degrees of freedom in their movement; and in EVR they used slightly more DoF than in the real-world task.

As a measure of task performance in body space, we use the Velocity Profile Error (VPE), as in Haar et al [21]. VPE is defined by the minimal correlation distance (one minus Pearson correlation coefficient) between the velocity profile of each joint in each trial to the velocity profiles of that joint in all successful trials. Like in the real-world, in the EVR environment we also found that VPE shows a clear pattern of decay over trials in an exponential learning curve for all joints (Figure 6A).

**Figure 6.**
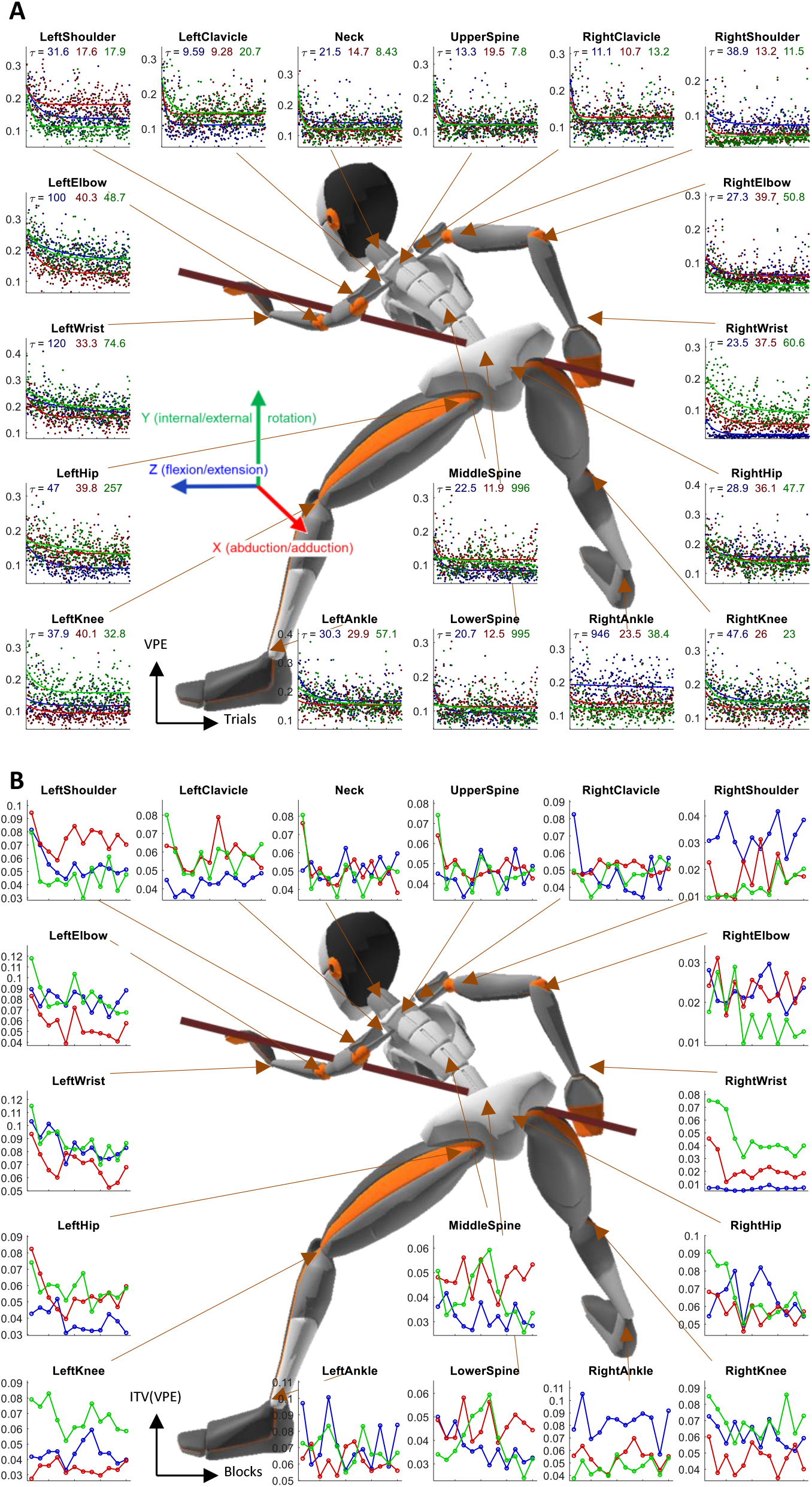
Learning over Joints. Velocity Profile Error (VPE) and Intertrial variability reduction across all joints in the EVR task. (**A**) The trial-by-trial VPE for all 3 DoF of all joints, averaged across all subjects, with an exponential fit. The time constants of the fits are reported under the title. Color coded for the DoF - blue: flexion/extension; red: abduction/adduction; green: internal/external rotation. (**B**) VPE intertrial variability (ITV) over blocks of 25 trials, averaged across all subjects. Data is averaged over the performance of 10 subjects in the novel EVR environment.

Intertrial variability in joint movement was also measured over the VPEs in each block. Unlike the real-world task, where learning was evident in the decay over learning of the VPE intertrial variability, in the EVR there was no such decay in most joints (Figure 6B). This is in line with the lack of decay in the intertrial variability of the directional error (Figure 3C&D).

## Discussion

In this paper, we present a novel embodied VR framework capable of providing controlled manipulations to better study naturalistic motor learning in a complex real-world setting. By interfacing real-world objects into the VR environment, we were able to provide the sense of embodiment to subjects’ experience in the real-world, which is missing in most VR environments. We demonstrate the similarities and differences between the learning in the EVR environment and learning in the real-world environment. Thus, we can now directly compare full-body motor learning in a real-world task between VR and the real-world. Before going into a detailed discussion of our findings, we reiterate the motivation for this work, namely, to develop a framework to allow neuroscience researchers to use our experimental techniques and protocols to perform in real-world tasks causal interventions by manipulating visual feedback. For example, changing the velocity of the virtual reality, manipulating the force perception of the shot, changing the ball directional error feedback to the subject, by modulating it at will or hiding the ball completely once it has been touched [e.g. 45]. In the context of rehabilitation, manipulations can purposefully increase task difficulty to enhance transfer [46].

There is considerable evidence that humans can learn motor skills in VR and transfer learning from the VR to the real-world. Yet, most of these evidences are in simplified movements [e.g. 47–50] while the successful transfer of complex skill learning remains a challenge. Accordingly, there is a real need to enhance transfer to make VR useful for rehabilitation and assistive technology applications [51, for review see 52]. Current thinking suggests that transfer should be enhanced the closer VR replicates the real-world [33,52]. The physical interactions in our EVR are generating accurate force, touch and proprioceptive perceptions which are lacking in conventional VR setups. This suggests that our EVR framework could thus enhance learning and transfer between VR and the real-world. A recent review on learning and transfer in VR [52] also highlights the fidelity and dimensionality of the virtual environment as key components that determine learning and transfer, which are also addressed in our setup. In that sense, EVR mock-up of real-life tasks, like the one presented here, should be the way forward as it addresses all the key components that determine learning and transfer in real-world tasks. Since both VR and motion tracking technologies are becoming affordable, EVR setups like the one presented here can be deployed in clinical rehabilitation settings in the foreseeable future. There is clearly much work to be done to validate these setups and to make them accessible for deployment (without programming expertise), but the use of affordable off-the-shelf hardware and open-source gaming engine makes it a relatively low-cost yet high-performance system.

### Limitations

The comparison of the ball trajectories between the EVR and the real-world environments highlight the similarity in the ball directions, which is the main parameter that determines task error and success. Nevertheless, there were significant velocity differences between the environments. These velocity differences were set to optimize subjects’ experience, accounting for deviations in ball physics due to friction, spin, and follow-through which were not modelled in the VR. Due to these deviations, in VR, the cue ball reaches its max velocity almost instantaneously while in the real-world there is an acceleration phase. For the current version of the setup, we neglected these differences, assuming it would not affect the sense of embodiment of very naïve pool players. Future studies, testing experts on the setup, would require more accurate game physics in the EVR.

Another limitation of the current EVR setup is that subjects are unable to see their own limbs in the environment, whereas in the real-world the positions of the subject’s own limbs may influence how the task is learned. We can probably neglect this difference due to the extensive literature suggesting that learning is optimized by an external focus of attention [for review see 53]. Thus, the lack of body vision should not significantly affect learning. The lack of an avatar (a veridical rendering of the person’s body) could also potentially affect VR embodiment. During a pool shot, most of the body is out of sight and only the hands holding the cue stick should be visible. While the hands and arms are not visible in our setup, the cue stick is, and thus there is a multi-sensory correlation between the haptic and proprioceptive feedback of handling the pool cue as a physical tool and the visual feedback from VR. This multi-sensory effect replicates the induction of embodiment strategy observed in e.g. the rubber hand illusion, as well as other experiments that measure embodiment through self-location [54,55].

Operating in VR requires acclimatisation and potentially learning a slightly altered visuomotor mapping (i.e. motor adaptation). While stereoscopic virtual reality technology seeks to present a true three-dimensional view of a scene to the user, distortions such as minification, shear distortion, or pincushion distortion can occur [56]. These distortions are dependent on the specific hardware setup, and an estimation of egocentric distances in VR may be off by up to 25% [56]. Moreover, stereoscopic displays induce a conflict between the different cues our visual system uses to infer depth and distance. Specifically, the mechanism of accommodation in the eye, which puts an object into focus, is also combined with other cues, such as vergence (movement of both eyes in opposite directions to maintain a single binocular vision) to estimate the object’s distance. This is an issue in VR as the focus point of all objects is on the screen immediately at nose-tip distance. This results in the accommodation-vergence conflict [57,58]. This conflict alters the visuomotor mapping and is a known cause of visual discomfort in VR [56,59,60]. Latest generation head-mounted displays (like the HTC Vive Pro used here) do address some of these issues, enable more accurate space perception than other head-mounted displays [61], and were validated and recommended for use in real navigation and even motor rehabilitation [62]. Yet, performing a real-world visuomotor task, such as a pool shot, probably requires some visuomotor learning of the VR environment, in addition to the motor learning of the task.

In the following, we compare motor learning between the VR setting and our physical setting [21] of our real-world pool task. To be clear, while in some scientific communities the term ‘learning’ is only used in the context of a retention test performed across multiple training sessions, we are taking here a human motor neuroscience view, where learning is the high-level term which includes short- and long-term motor learning phenomena [for review see 63, 64]. Thus, we use the term ‘learning’ here to describe changes in task performance within a session. Motor learning in the EVR task and the physical real-world task [21] showed many similarities but also intriguing differences. The main trends over learning that were found in the physical real-world task include the decrease in directional error, decrease in directional intertrial variability, decrease in shoulder movement and increase in elbow rotation, decrease in joints VPE, and decrease in joints VPE intertrial variability [21]. In the EVR environment, we found the same general trends for all these metrics, except for those of the intertrial variability. We do however see a systematic difference in learning rates between VR and real-world when comparing the directional error and VPE trends. Across the board, we see less learning in VR compared to the physical real-world. This is presumably due to the additional learning of the altered visuomotor mapping required in VR.

The decay of intertrial variability over trials is a prominent feature of skill learning [64–71], but not found in motor adaptation experiments. In the physical pool paradigm, we found two types of subjects that differ in their intertrial variability decay as well as other behavioural and neural markers [22]. Altogether, this suggested the contribution of two different learning mechanisms to the task: error-based adaptation and reward-based reinforcement learning, where the predominant learning mechanism is different between subjects. Here, the lack of intertrial variability decay and the overall differences in the learning curve between the group that learned the task in the EVR setting and the group that learned it in the physical real-world setting, suggests potential differences in the learning mechanisms used by the subjects who learned the task in the EVR. Presumably, in the EVR environment, the predominant learning mechanism was error-based adaptation. This may be attributed to the fact that all subjects were completely naïve to the EVR environment and had to learn not just the billiards task but also how to operate in EVR.

Like in the physical task, in the EVR setting, we also found motor learning to be a holistic process – all body joints are affected as a whole by learning the task. This was evident in the decrease in the velocity profile error (VPE) and the intertrial variability over learning (Figure 6). As we highlight in [21], this holistic aspect of motor learning is known in sport science [e.g. 72–74] but is rarely studied in motor neuroscience and motor rehabilitation, which are usually using simplistic artificial tasks and measure movement over only one or two arm joints. This congruence between physical and EVR setting suggests that while there are clear differences between the environments in the learning and performance in the task level (as discussed above), there are strong similarities in the body level [75] which can potentially enhance transfer.

### Conclusions

In this study, we have developed an embodied VR framework capable of applying visual feedback manipulations for a naturalistic free-moving real-world skill task. We have also demonstrated the similarities in learning progression for a pool billiards shot between the EVR and the real-world and have confirmed real-world findings that motor learning is a holistic process which involves the entire body from head to toe. By manipulating the visual feedback in the EVR we can now further investigate the relationships between the distinct learning strategies employed by humans for this real-world motor skill. Our approach is useful for rehabilitation, as it overcomes the limit to VR immersion, one of the reasons why many VR rehabilitation approaches are based on 3D graphics viewed on computer monitors and not actual VR using head-mounted displays. Our work highlights the potential of real world-tasks being used in VR in rehabilitation, but also in VR based skill training such as biosurgery.

## Authors’ contributions

SH and AAF conceived and designed the study; SH, GS and AAF developed the experimental setup; GS acquired the data; GS and SH analysed the data; SH, GS and AAF interpreted the data; SH drafted the paper; SH and AAF revised the paper

## Acknowledgements

We thank our participants for taking part in the study. The study was enabled by financial support to a Royal Society-Kohn International Fellowship (NF170650; SH & AAF).

